# Automated, modular assembly of reconstituted cell-free systems from *in vitro*-produced components

**DOI:** 10.64898/2025.12.23.696326

**Authors:** Byeongmin Chae, Hyeongwoo Park, Hyewon Jeon, Jihoo Lee, Haneul Jin, Jiwoon Jeong, Jinjoo Han, Jeong Wook Lee, Joongoo Lee

## Abstract

High-performance cell-free protein synthesis has transformative potential for synthetic biology, yet the prohibitive costs of PURE kits and the labor intensity of in-house preparation have restricted accessibility and scalability. We developed *i*-POPFLEX (Purified components OPtimized for FLEXible protein expression), a modular cell-free protein synthesis system in which 34 translation factors, individually synthesized *in vitro*, are assembled using automated liquid handling. This workflow minimizes manual input and supports parallelized production, generating complete, ready-to-use systems within two days. *i*-POPFLEX achieves up to 5-fold higher protein yields and a 95 % cost reduction (20-fold lower cost) compared with commercial kits. Its flexible architecture also enables selective component inclusion for genetic code reprogramming and site-specific incorporation of non-canonical amino acids. By coupling modular design with automation, *i*-POPFLEX provides an accessible, customizable, and economically viable platform for next-generation biomanufacturing workflows.

**Technology readiness:** *i*-POPFLEX is a reconstituted cell-free system (CFS) composed of 34 translational components that are individually synthesized *in vitro* and assembled by automation. This system achieves up to 5-fold higher protein yields with a 95 % cost reduction (20-fold lower; <USD 0.06 µL^-1^) compared with commercial PURE kits (USD 1.36 µL^-1^) and reduces preparation time from four days to two. The workflow has been validated across diverse protein and peptide synthesis applications, including genetic code reprogramming and site-specific incorporation of non-canonical amino acids, and integrates seamlessly with benchtop automated liquid-handling platforms. These capabilities place *i*-POPFLEX at Technology Readiness Level (TRL) 4-5: integrated components validated in a laboratory setting and advancing toward pilot-scale readiness.

For full-scale deployment, several key challenges remain. Matching crude lysate costs (USD 0.03 µL^-1^) will require bulk reagent production, optimized procurement strategies, and specialized equipment for high-throughput component synthesis. Enabling milliliter- and liter-scale production will demand improvements in upstream synthesis, downstream purification, and quality control to ensure consistent performance at industrial throughput. Overcoming these barriers would enable production of thousands to millions of fully assembled *i*-POPFLEX systems for biofoundry-scale enzyme engineering, metabolic pathway prototyping, and large-scale variant screening, democratizing reconstituted CFS technology, and accelerating adoption in next-generation biomanufacturing pipelines.

## Introduction

**Cell-free systems (CFSs)** (see **Glossary**) enable gene expression and protein synthesis outside living cells in programmable biochemical environments, offering exceptional flexibility for synthetic biology and industrial biomanufacturing [1–5]. CFSs now drive bio-based innovations across diverse domains, including (i) rapid prototyping of metabolic pathways [6,7] and bottom-up construction of synthetic cells [8,9]; (ii) high-yield expression of proteins that are toxic or unstable *in vivo* [6,10–12]; (iii) on-demand and on-site diagnostics and biomolecule sensing [13,14]; (iv) accelerated enzyme discovery using high-throughput screening [15,16]; and (v) site-specific incorporation of **non-canonical amino acids (ncAAs)** to produce advanced antibody-drug conjugates (ADCs) [17–19].

CFSs generally fall into two categories: crude-lysate and reconstituted systems assembled from individually purified factors. Crude lysates retain native biochemical properties of ribosomes, **tRNAs**, metabolic enzymes, and cofactors (20-40 % of the *E. coli* proteome) [20–23], supporting milligram- to gram-scale protein yields at low cost (∼USD 0.03 µL^-1^) from microliters (µL) to industrial multi kiloliter volumes (up to 4.5 kL) [24–26]. However, the undefined composition of crude lysates limits reproducibility, systematic optimization, and genetic code reprogramming [27]. In contrast, reconstituted systems, exemplified by the PURE (Protein synthesis Using Recombinant Elements) platform [28] and commercial kits such as PURExpress^TM^ and PUREfrex^®^ 2.0 [29–32], offer a fully defined and interference-free (e.g., nuclease- and protease-free) environment with exceptional reproducibility, making them ideal for quantitative biochemistry [33], genetic circuit prototyping [34], and mechanistic studies of enzymes [35]. Yet, high reagent cost (USD 1.00-1.36 µL^-1^; [36,37]) limits their widespread use in automated biofoundries, where large libraries of enzyme variants must be expressed and screened economically. For example, screening a modest library with five randomized positions of a protein (20^5^ = ∼10^6^ variants) would cost ∼USD 1,000 even at nanoliter reaction volumes, and scaling to a 10-site library (20^10^ = ∼10^13^ variants) could exceed USD 10 billion.

To overcome these barriers, Lavickova et al. developed a One-Pot PURE method [36] combining co-culturing with single-step purification, reducing costs to USD 0.09 µL^-1^, while Zhang et al. further improved system robustness [37]. Nevertheless, manual strain engineering and purification for each component remains labor-intensive, time-consuming, and susceptible to batch variability. These factors would limit the speed, reliability, and reproducibility required for modern biofoundry operations.

Here, we introduce a semi-automated workflow for producing a reconstituted CFS that increases synthetic capability by up to 5-fold (**Figure S1** in the Supplemental Information online), lowers reagent costs by as much as 95 % (**Table S1** in the Supplemental Information online), and halves both preparation time and labor (**Table S2 and S3** in the Supplemental Information online), supporting efficiency and cost reduction aligned with modern Design-Build-Test-Learn cycles in synthetic biology workflows. We first show that our approach matches or exceeds the performance of previously reported reconstituted CFSs prepared by manual purification when using an automated chromatography platform and alternative in-house reagents. We next demonstrate that all translational factors, except for **T7 RNA polymerase (T7RNAP)**, are produced entirely *in vitro,* thereby decoupling the protein synthesis step from living materials, and assembled on an automated platform with high precision and speed.

Moreover, we show that the workflow supports on-demand customization of component sets, enabling ncAA incorporation via sense codon reassignment (**Figure 1**).

**Figure 1.**
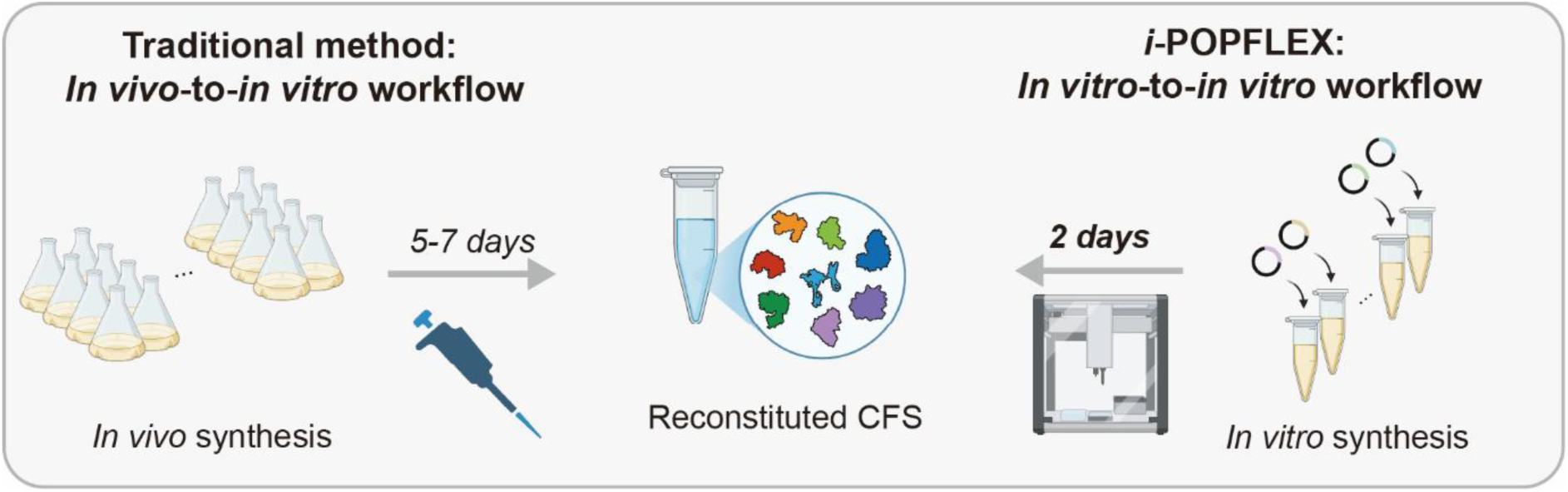
*i*-POPFLEX: A modular reconstituted CFS containing in-vitro produced translational machinery. In contrast to traditional *in vivo* expression and manual purification, our lysate-based, fully *in vitro* approach synthesizes all components and assembles them with high precision using an automatic liquid-handling platform.

Taken together, these advances bridge the gap between laboratory-scale innovation and practical implementation, transforming reconstituted CFSs from specialized experimental tools into robust, higher-capacity platforms for next-generation biofoundry application and the manufacture of high-value biomaterials.

## Results

### Challenges in replicating *in vivo* synthesis of translational machinery

While the reconstituted CFS offers substantial potential for cost-effective protein synthesis and high-throughput enzyme analysis in biofoundries, reproducibility across individual laboratories remains a challenge. To address this, we first attempted to replicate the One-Pot PURE system, which relies on co-culture and co-purification of all protein components required for *in vitro* translation [36,38]. Following previous studies, we employed two bacterial strains: an *E coli* BL21 (DE3) derivative (T7 Express Competent *E. coli*, NEB) and DH5-α (NEB^®^ 5-alpha Competent *E. coli*, NEB) capable of recognizing T7 and T5 promoters, respectively. Because the optimized M15 strain used in earlier work [36] was not commercially available, we substituted DH5-α for T5-driven expression, which relies on native σ70 RNA polymerase to recognize the T5 promoter. Genes encoding the protein translational machinery, each containing a His-tag, were cloned into T7-based pET and T5-based pQE plasmids. Cells were cultured and lysed, and the His-tagged proteins were purified according to previously published protocols [38]. Despite our efforts, our preparation did not show activity in protein synthesis such as superfolder green fluorescent protein (sfGFP) expression. Although the DH5-α strain is not optimal for protein expression, we observed inconsistent growth among strains harboring pQE plasmids, consistent with previously reported batch-to-batch variability of the One-Pot PURE system [39]. For instance, Ngo et al. reported deletions in the T5 promoter region of pQE plasmids after prolonged storage at −80 °C, while the Murray group observed reduced protein expression efficiency in dual-strain systems preserved in glycerol-containing solutions [37]. To address these limitations and improve the reliability and efficiency of protein expression, we shifted our focus to replicating the single-strain One-Pot PURE (ssOP) system developed by the Murray group [37].

To ensure uniform growth rates across all expression strains, we replaced the original pQE vectors with T7 promoter-based pET vectors and subcloned 14 out of the 16 expression plasmids (**Table S4 and S5** in the Supplemental Information online) into the *E. coli* BL21 (DE3) derivative strain, excluding those for T7RNAP and release factor 1 (RF1). Previous studies have shown that expressing T7RNAP from a T7 promoter in mixed cultures induces an autocatalytic feedback loop, resulting in severe overexpression and cell death [37]. To mitigate this instability, we supplied T7RNAP externally as a commercial preparation (NEB), ensuring efficient transcription without compromising culture viability. Additionally, we intentionally omitted RF1, a key translation termination component, to enable flexible reassignment of the UAG (amber) codon for the incorporation of ncAAs. Without RF1, the UAG codon, normally a stop signal, can be repurposed to encode a variety of ncAAs, thereby expanding the versatility of our reconstituted system.

Of the 36 individual components required for protein translation, the 34 corresponding pET vectors were separately transformed into *E. coli*. Following transformation, all cultures were grown individually in LB medium overnight. The cultures were then pooled into 500 mL of fresh LB medium and grown to saturation. During this step, **EF-Tu** was supplied in excess (30:1, v/v) due to its critical role in translation as insufficient EF-Tu levels can lead to poor tRNA delivery, ribosome stalling, and reduced protein yields [36,40]. Maintaining abundant EF-Tu ensured that the *in vitro* translation system was not limited during protein synthesis. After culturing, the cells were lysed via sonication, centrifuged, and the His-tagged proteins were purified from supernatant. To improve upon the conventional gravity-flow purification used in previous ssOP strategies, we implemented an automated fast protein liquid chromatography (FPLC) system (**Figure 2A**). Adoption of FPLC reduced protein elution times from several hours to ∼20 minutes, while also increasing protein yield, purity, and reproducibility, all of which are critical for assembling a robust reconstituted CFS. This optimized workflow shortened the overall procedure to 3-4 days, compared to the 5-7 days required for conventional manual assembly (**Figure S2 and Table S2** in the Supplemental Information online).

**Figure 2.**
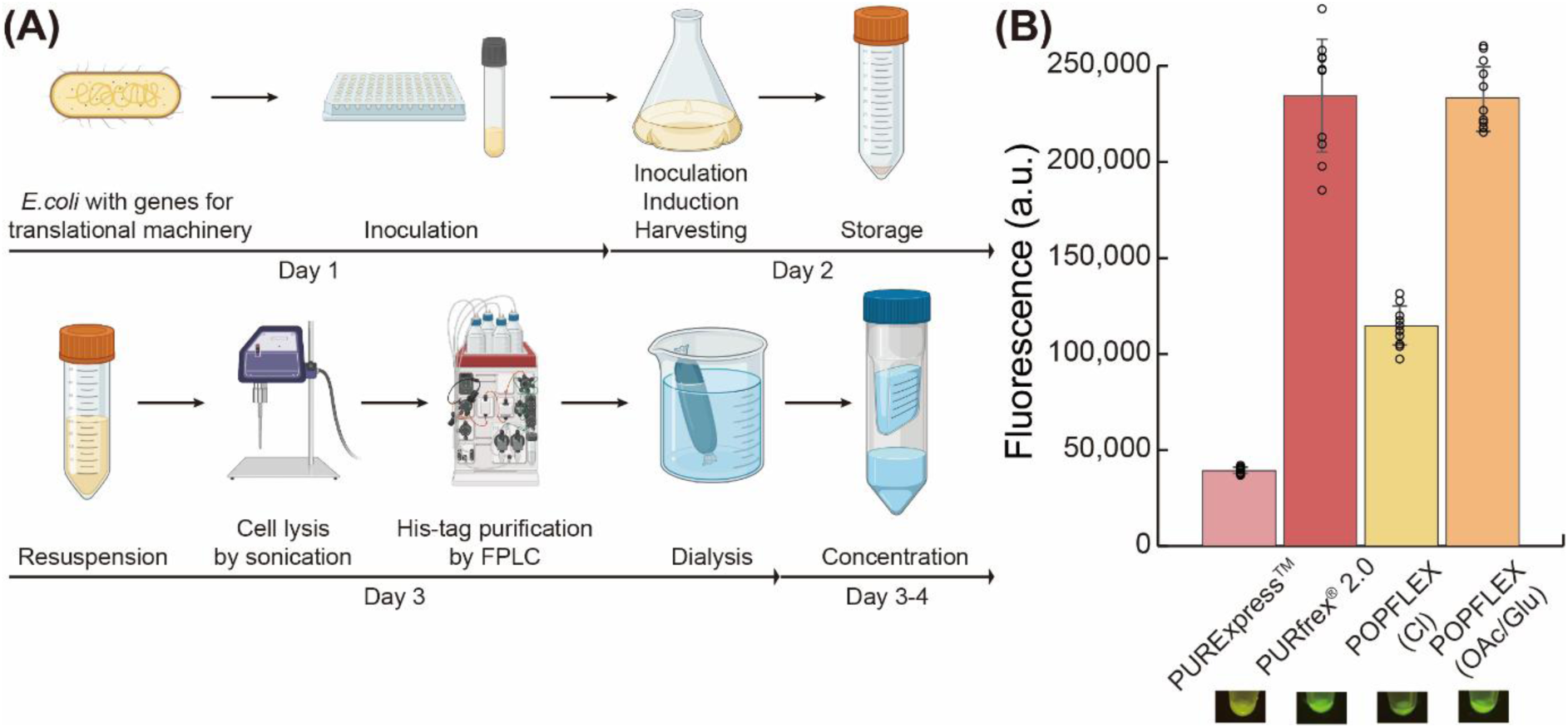
Reconstituted cell-free system using automated liquid chromatography equipment. (**A**) Schematic overview of POPFLEX preparation. In contrast to conventional methods, FPLC enabled fast and highly reproducible protein purification. (**B**) Comparison of sfGFP fluorescence from different system configurations. Replacing chloride-based salts (yellow bar) with acetate- and glutamate-based alternatives improved POPFLEX protein expression by 2-fold (orange bar), reaching levels comparable to the commercially available PURExpress^TM^ (NEB, pink bar) PUREfrex^®^ 2.0 system (GeneFrontier, orange bar). Error bars represent the standard deviation of *n* = 10 independent experiments.

Using the FPLC-purified protein mixture, we first evaluated its capacity for protein synthesis. Following protocols established in previous studies [36,37], the purified protein mixture was concentrated and adjusted to a final concentration of 1 mg·mL^-1^, then combined with buffers containing commercial *E. coli* endogenous tRNA and purified ribosome (GeneFrontier), and a DNA template encoding sfGFP. Initial tests revealed lower sfGFP expression levels (yellow bar, **Figure 2B**) compared to the commercial PURExpress^TM^ and PUREfrex^®^ 2.0 kits (pink and red bar, respectively, **Figure 2B**). To probe this discrepancy, we investigated the influence of chloride ions originating from MgCl_2_ and KCl present in conventional dialysis buffer. While chloride-based are essential cofactors in transcription and translation reactions, chloride ions have been reported to inhibit enzymatic activities, particularly reducing the catalytic efficiency of T7RNAP [41,42]. To test an alternative ionic environment, we substituted MgCl_2_ and KCl with acetate- and glutamate-based salts (Mg(OAc)_2_ and KGlu, respectively). This substitution led to a 2-fold increase in sfGFP fluorescence (orange bar, **Figure 2B**), indicating enhanced transcriptional and translational activity. Taken together, our optimized ssOP system, which integrates FPLC-based His-tag purification and physiologically relevant acetate/glutamate buffering significantly enhances both the stability and productivity of reconstituted CFS workflows for *in vitro* protein synthesis. We refer to this optimized system as Purified components OPtimized for FLEXible protein expression (POPFLEX), a modular platform designed for high yield cell-free protein synthesis system. Our POPFLEX protocols typically yield 250 µL of purified protein solution at 10-12 mg·mL^-1^ from 500 mL of culture. This output is sufficient for 50 cell-free reactions at a 25 µL scale, representing up to a 5-fold improvement in productivity and reagent utilization compared to conventional approaches. Notably, the standard PURExpress^TM^ kit (E6800S) supplies reagents for only 10 reactions of the same volume.

### Shift to lysate-based cell-free platforms for translation machinery production

Traditional approaches to constructing reconstituted CFSs have relied on an *in vivo*-to-*in vitro* workflow, in which translation factors are produced within living cells, then purified and assembled *in vitro*. However, the preparation of translational machinery *in vivo* remains a major bottleneck, significantly limiting the efficiency of CFS assembly. From culture to final protein mixture preparation, our POPFLEX protocols typically require 4 days (**Figure 2A**). These labor-intensive and time-consuming steps limit scalability and throughput. To overcome these constraints, we shifted toward a fully *in vitro* strategy for producing the 34 translational components. This approach offers substantial advantages: the open nature of CFSs enables rapid, parallelized, automation-compatible production, reduces dependence on large-scale cell culture, and lowers time and cost requirements. By integrating automated liquid handling, this strategy lays the groundwork for efficient and economically sustainable pipelines in high-throughput synthetic biology and enzyme engineering.

To implement this strategy, we produced all POPFLEX components using lysate-based CFSs. We selected the *E. coli* cell lysate platform for its robustness, low cost, and broad applicability, including the efficient production of proteins with diverse non-canonical amino acids (ncAAs) [29,31,43–46]. Briefly, lysates were prepared from 1 L cultures by sonication and clarified by centrifugation to remove debris. Each translation component was cloned into pJL1 expression plasmids, which incorporate optimized promoter spacing, ribosome binding sites, and flanking sequences around the T7 promoter to enhance transcription and translation efficiency in CFSs [47,48]. The pJL1 vector was particularly advantageous for its low background expression and favorable transcription-translation dynamics, supporting protein yields up to the gram-per-liter (g·L^-1^) scale [1,49,50].

To produce the 34 translational components used in POPFLEX, we carried out 45 µL lysate-based cell-free reactions. Each reaction contained *E. coli* S30 cell lysate, in-house prepared Solution A and B, and a pJL1 plasmid encoding a single translational component. To verify expression, we manually purified the *in vitro* synthesized proteins using cobalt agarose beads and analyzed them by SDS-PAGE. All proteins were successfully expressed (**Figure 3A-D**), except release factor 2 (RF2, **Figure S3** in the Supplemental Information online). We suspected that RF2 expression was impaired due to stable secondary structure in its T7-scribed mRNA that hindered ribosome progression. This likely impeded translation initiation, leading to misfolded or absent protein [51,52]. To address this, we substituted the circular plasmid with a linear DNA template for RF2. Because linear DNA templates are susceptible to **RecBCD**- mediated degradation in *E. coli* cell lysates [53], we flanked the template with 200-bp non-coding sequences to improve stability against enzymatic degradation [54,55]. As hypothesized, substituting the circular plasmid with a linear DNA template successfully enabled RF2 expression (Lane 2, **Figure 3C**).

**Figure 3.**
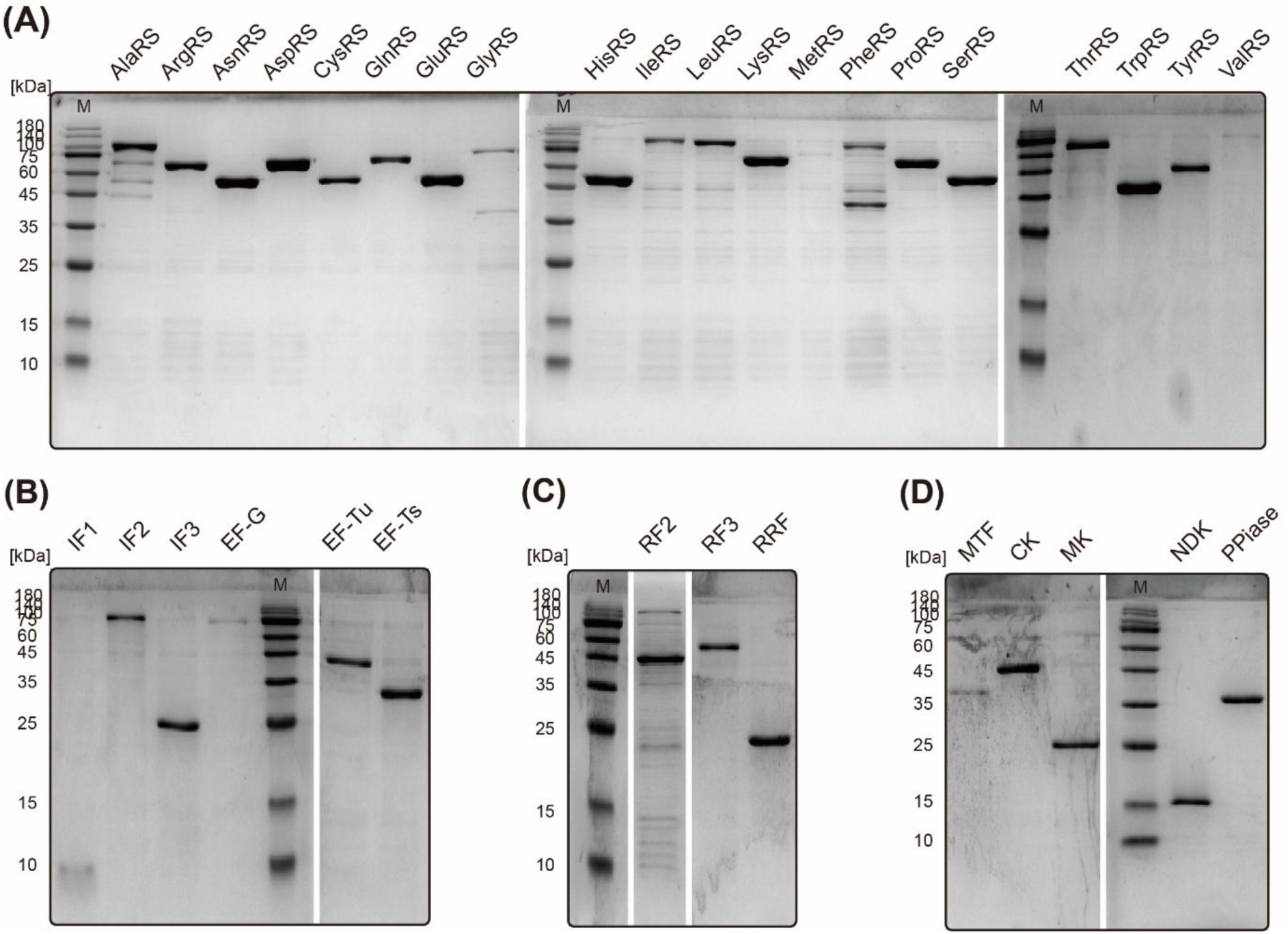
SDS-PAGE analysis of *in vitro*-synthesized and purified translational machinery components. (**A**) 20 aminoacyl-tRNA synthetases (aaRSs). (**B**) Initiation and elongation factors (IEFs). (**C**) Release factors (RFs); RF1 was omitted in *i*-POPFLEX system to enable reassignment of the amber codon (UAG) for ncAA incorporation. RF2 was synthesized using a linear DNA template (Lane 2). (**D**) Energy regeneration factors (ERFs). The concentration of EF-Tu (Lane 6), as determined by Bradford Assay, was 1.98 mg·mL^-1^. The lowest detectable band intensity for MetRS or ValRS on the gel corresponded to ∼0.4 mg·mL^-1^, based on densitometric analysis using ImageJ (**Table S6** in the Supplemental Information online). These results indicate that our *i*-POPFLEX system is functional when each protein is present at a minimum of 0.4 mg·mL^-1^, and at a final combined concentration of roughly 5-7 mg·mL^-1^ after concentration with a centrifugal filter. M: molecular marker.

We also attempted to produce T7RNAP using this lysate-based approach, even though expression in the pJL1 vector system was poor [56]. However, repeated attempts to clone T7RNAP into T7 promoter-driven vectors (e.g., pET21a and pJL1) consistently yielded plasmids with incorrect amino acid sequences. Specifically, we observed deletion of two residues (‘FA’: F894 and A895 in our construct, including the N-terminal His-tag and linker; equivalent to F882 and A883 in the native protein) within the C-terminal FAFA motif, a critical sequence for maintaining enzymatic activity in the ‘foot’ domain [57]. This observation is consistent with previous reports that pJL1-T7RNAP constructs frequently accumulates mutations [56].

Although SDS-PAGE analysis confirmed that all 34 translational components, including RF2, migrated at their expected molecular weights, our initial attempts to synthesize sfGFP using this *in vitro*-produced protein mixture were unsuccessful (**Figure 4**). Several reactions produced cloudy precipitates after overnight incubation, which led us to hypothesize that additional sequences present in circular DNA templates, such as antibiotic resistance genes and replication origins, might introduce transcriptional interference *in vitro*, potentially contributing to protein misfolding or aggregation.

**Figure 4.**
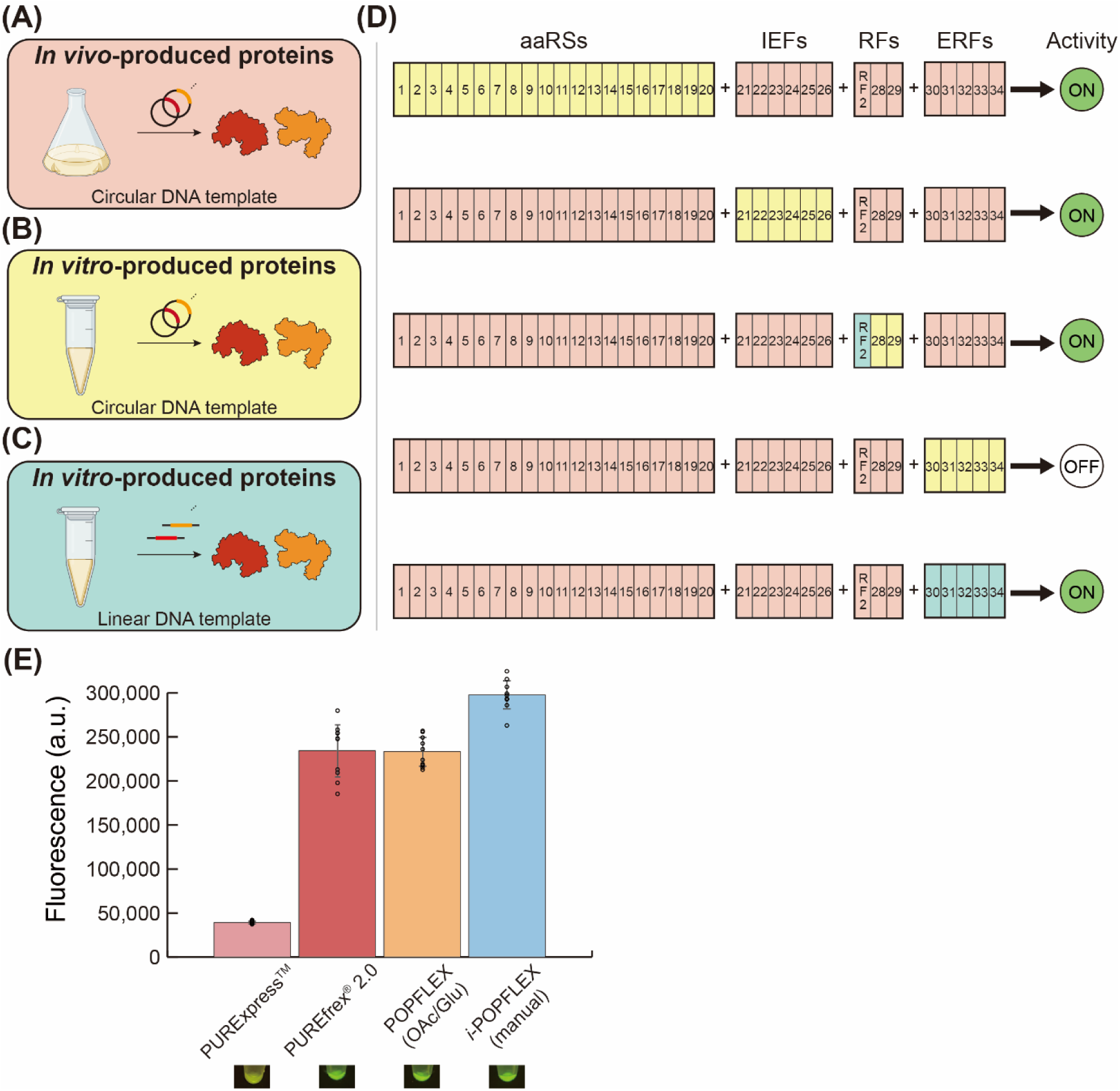
Linear DNA templates are required for efficient *in vitro* synthesis of certain translational components. Initial attempts to synthesize sfGFP using all 34 components produced in S30 lysate (Figure 3) were unsuccessful. To identify the problematic components, we grouped them into four categories; aaRSs, IEFs, RFs, and ERFs. Using *in vivo*-produced components capable of expressing sfGFP (**A**), we systematically designed four test sets, each containing one group produced *in vitro* (**B**). This analysis revealed that the set containing *in vitro*-produced ERFs was unable to produce sfGFP (**D**). Replacing circular DNA templates with linear templates (**C**) for ERF expression restored sfGFP synthesis, confirming the requirement of linear DNA for efficient RF2 and ERF production (**D**). (**E**) Fluorescence measurements showed that sfGFP produced using the *i*-POPFLEX system exhibited higher signal intensity compared to POPFLEX and commercially available reconstituted systems (PURExpress^TM^ and PUREfrex^®^ 2.0). Error bars represent the standard deviation of *n* = 10 independent experiments.

To identify functionally inactive components, we categorized the 34 proteins into four functional groups: 20 **aaRSs**, 6 IEFs, 3 RFs, and 5 ERFs. We then systematically tested each group by expressing it *in vitro*, while supplying the remaining components from *in vivo*-prepared proteins for POPFLEX (**Figure 4A-C**). For example, we reconstituted the system with *in vitro*-produced ERFs and *in vivo*-produced aaRSs, IEFs, and RFs. Failure to express sfGFP would indicate inactivity of the *in vitro*-produced ERFs (**Figure 4D**). RF2 was included using the linear DNA template in the ERF mixture.

This process was repeated for all four groups. The results revealed that sfGFP expression failed only when the ERFs were supplied from *in vitro* production, suggesting these proteins were inactive due to the same secondary structure issue observed with RF2. To resolve this, we synthesized linear DNA templates for the five ERF components (**Figure 4C**), flanked by 200 bp non-coding sequences at both ends to protect against exonuclease degradation, as described earlier. Using these linear templates, we re-expressed the ERF proteins *in vitro*, and with all 34 components, now including the ERFs, we successfully achieved sfGFP expression, confirming the full reconstitution of an active cell-free protein synthesis system (**Figure 4D**).

To assess the activity of the *in vitro*-produced components, we compared their protein synthesis performance with the original POPFLEX system and two commercially available reconstituted systems (PURExpress^TM^ and PUREfrex^®^ 2.0). Notably, the *i*-POPFLEX system (blue bar, **Figure 4E**), comprising *in vitro*-produced POPLEX components, exhibited higher fluorescence of sfGFP and reproducibility (*n* = 10) compared to both reference systems.

### Semi-automation of translational machinery preparation for a reconstituted CFS

While effective, manual handling 34 separate cell-free reactions was labor-intensive (∼2-3 h) and error-prone, with small deviations potentially compromising overall performance of the *i*-POPFLEX. To address these limitations, we integrated the workflow into an automated liquid-handling platform (Opentrons OT-2), enabling standardized *in vitro* synthesis of all 34 translational components (**Figure 5A**).

**Figure 5.**
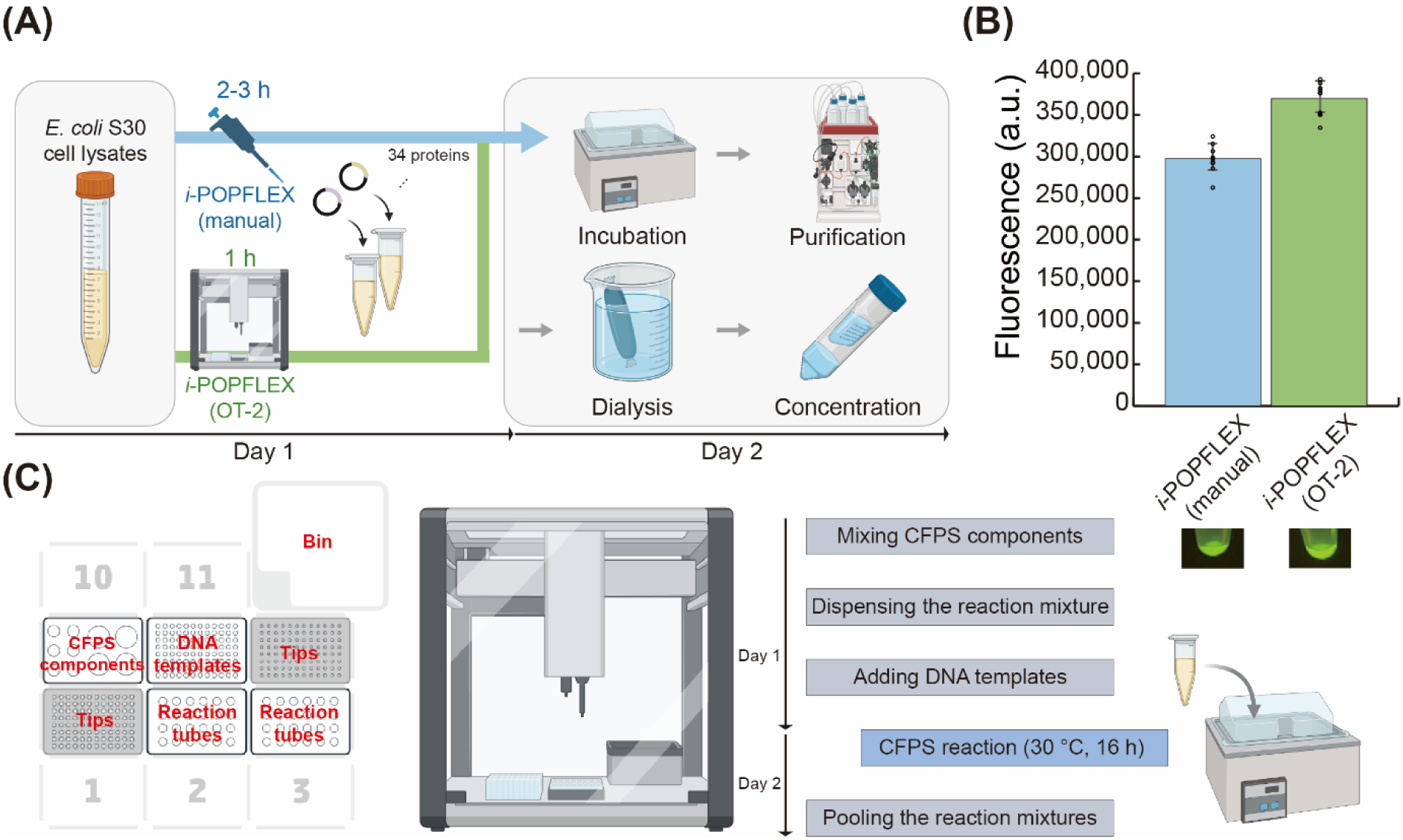
Semi-automated *i*-POPFLEX platform for synthesis of translational machinery. (**A**) Schematic overview of *i*-POPFLEX preparation. All manual mixing steps were executed on an automated liquid-handling platforms (OT-2), enabling efficient and modular assembly of reconstituted CFSs. (**B**) sfGFP produced using automated *i*-POPFLEX shows higher fluorescence compared to manually assembled *i*-POPFLEX. Error bars represent the standard deviation of *n* = 10 independent experiments. (**C**) OT-2 workflow and representative deck layouts. Four steps were carried out within the OT-2 chassis, with incubation performed externally (blue highlighted box).

The OT-2 workflow comprises five key steps; i) preparation of multiple S30 lysate-based cell-free reactions, ii) automated distribution of DNA templates into individual microtubes, iii) incubation under controlled conditions, iv) pooling of 34 individual reactions into a single collection tube, and v) purification of the combined protein components (**Movie S1 and S2** in the Supplemental Information online). Steps i), ii), and iv) were executed within the OT-2 chassis, while incubation was carried out in a water bath and the pooled sample was manually injected to an automated chromatography system for final purification.

The purified protein mixture was dialyzed and concentrated using a centrifugal protein concentrator (MWCO 3 kDa, Amicon) to a final concentration of 5-7 mg·mL^-1^. The concentrated proteins were then combined with in-house-prepared buffers (*E. coli* endogenous tRNAs + 4X energy solution), purified ribosome (GeneFrontier), T7RNAP, and the sfGFP DNA template to initiate the reaction. Following incubation at 37 °C for 3 h, fluorescence was measured to assess protein synthesis capacity relative to the manually prepared reconstituted system. *i*-POPFLEX assembled on the OT-2 platform achieved a ∼20 % increase in sfGFP expression levels (green bar, **Figure 5B**) compared with manually prepared *i*-POPFLEX (blue bar, **Figure 5B**). The total manufacturing time on the semi-automated platform was 1 h, representing a 2-3 fold reduction compared with manual preparation. Together, these results showed that the semi-automated workflow provides an efficient foundation designed for future scalability in generating in-house reconstituted CFSs, with *i*-POPFLEX achieving up to a 9.4-fold improvement in protein synthesis performance over commercially available PURE kits.

### Tailored *i*-POPFLEX systems enable efficient ncAA incorporation and synthesis of bioconjugation-ready biomaterials

The automated pipeline of *i*-POPFLEX allows precise control over individual components in reconstituted CFSs by preparing a DNA template, expressing an enzyme, and co-purifying components. This modular architecture allows selective omission of endogenous components, such as RFs or aaRSs, which can act as competitors for ncAA incorporation by utilizing their cognate codons. Removing these competitors permits both nonsense and sense codon reassignment on demand.

To demonstrate this flexibility, we designed two customized *i*-POPFLEX systems, one lacking RF1 and another containing a reduced set of aaRS, both including the full set of initiation, elongation, release, and other factors required for translation (**Table S7** in the Supplemental Information online). We built minimal DNA templates encoding Strep-tag derivatives (WSHPQFEK and MWHPQFEK, respectively) under T7 promoter control to serve as reporter peptides. The first system, lacking RF1, was used to incorporate ncAAs at the N-terminus (**X-**WSHPQFEK) via the nonsense (UAG) codon. The second system contained only eight of the 20 aaRSs (MetRS, TrpRS, HisRS, ProRS, GlnRS, PheRS, GluRS, and LysRS) required for the reporter peptide synthesis, enabling ncAA incorporation at the C-terminus (MWHPQFEK-**Y**) through sense codons. By removing serine from the reporter peptide and reducing tRNA competition, this design expanded the genetic code through sense codon reassignment and minimized serine misincorporation at the ACC codon caused by wobble base pairing [58], thereby improving ncAA incorporation efficiency.

Site-specific ncAA incorporation requires tRNAs bearing anticodons for the target to be acylated with the ncAA. Traditionally, this acylation relies on engineering aaRS to accept ncAAs, a technically demanding process often that often yields limited substrate promiscuity [58,59]. Instead, we employed **flexizyme (Fx)**, ribozymes capable of acylating any CCA-ending tRNA with diverse ncAAs by recognizing an aromatic leaving group, enabling rapid charging of structurally diverse substrates. Recent improvements in Fx-substrate design rules have further expanded their applicability [31,46].

We synthesized two Fx-compatible substrates (substrates (**1**) and (**2**): an azido-lysine analog and a propargyl-glutamate analog, respectively; see **Figure 6** and the Supplemental Information online) chosen for their orthogonal reactivity in **Cu(I)-catalyzed azide-alkyne cycloaddition (CuAAC)**, a widely used ligation reaction in ADCs, cell-surface labeling, biomaterial fabrication, and intracellular imaging [17,60–62]. Each ncAA was designed with an extended side chain to spatially separate the reactive group from the protein surface, facilitating conjugation efficiency [63]. Briefly, substrate (**1**) was prepared via diazotransfer to introduce the azide, dinitrobenzyl (DNB)-group attachment, and global deprotection. Substrate (**2**) was synthesized by attaching a DNB-group, followed by deprotection/re-protection, propargyl esterification, and final Boc removal (see Supplemental Information online).

**Figure 6.**
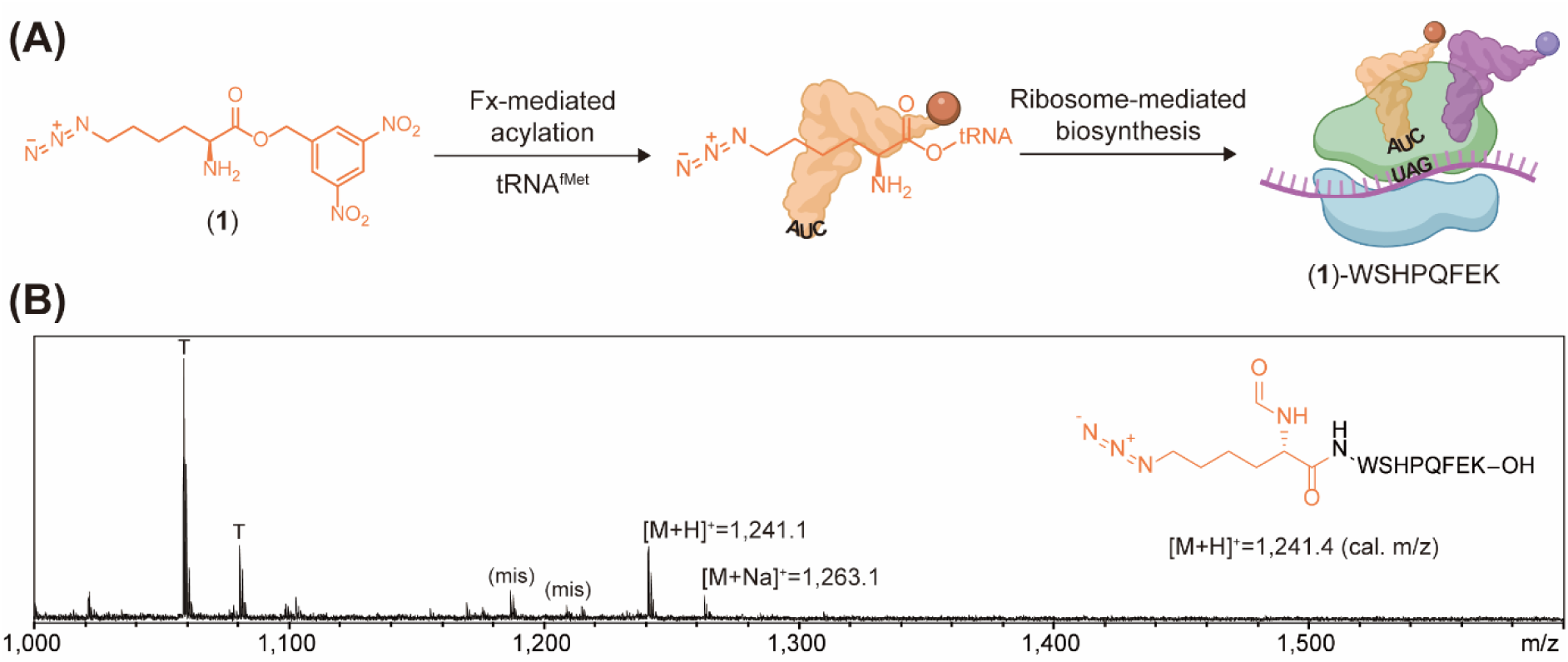
*i*-POPFLEX supports non-AUG translation initiation by omitting RF1. (**A**) Substrate (**1**) was activated with a DNB ester and charged onto tRNA^fMet^(CUA) by Fx. The charged (**1**):tRNA conjugate was then added to *i*-POPFLEX reaction containing DNA template, enabling the N-terminal incorporation of substrate (**1**). (**B**) MALDI-TOF mass spectra of the resulting peptides. Peaks with **T** indicate a truncated peptide lacks the first one ncAA at the N-terminus (WSHPQFEK). Peaks marked as **(mis)** indicate peptide containing a misincorporated Met at the **X** residue. Mass spectra are representative of *n* = 3 independent experiments. The theoretical masses of target peptide ((**1**)-WSHPQFEK) are [M+H]^+^ = 1,241.4 (cal) and [M+Na]^+^ = 1,263.4 (cal).

To optimize acylation efficiency, we monitored Fx-mediated charging of these substrates onto a tRNA mimic (microhelix) at two pH values using three different Fxs.

After determining optimal acylation conditions (**Figure S4** in the Supplemental Information online), we charged substrate (**1**) onto tRNA^fMet^(CUA), transcribed *in vitro* by T7RNAP. The charged tRNAs were purified and added to the *i*-POPFLEX reaction in which the reporter’s AUG initiation codon was replaced with UAG. This system produced (**1**)**-**WSHPQFEK (**Figure 6A**). Streptavidin affinity purification followed by MALDI-TOF analysis showed incorporation of substrate (**1**) at the N-terminus of the target peptide ([M+H]^+^ = 1,241.1 (obs)) (**Figure 6B**). Although truncated peptides (denoted as T) lacking substrate (**1**) were produced as a major product, this result is important because it demonstrates that a stop codon can serve as an alternative initiation codon without additional translational machinery engineering [64].

To further demonstrate sense codon reassignment, we charged substrates (**1**) and (**2**) onto tRNA^Pro1E2^(GGU) and tRNA^Pro1E2^(GAU), respectively, under the optimized Fx condition (**Figure 7A**). The purified tRNAs were added to *i*-POPFLEX reactions containing reporter constructs in which the codons ACC and AUC (decoded as Thr and Ile in cells) were placed at the C-terminus, yielding M-WHPQFEK-(**1**) and M-WHPQFEK-(**2**), respectively. MALDI-TOF confirmed successful site-specific incorporation of (**1**) ([M+H]^+^ = 1,285.0 (obs)) and (**2**) ([M+H]^+^ = 1,298.0 (obs), **Figure 7B,C**). Subsequent CuAAC with complementary azide or alkyne probes produced triazole-linked conjugates of which masses matched theoretical values ([M+H]^+^ = 1,341.1 and 1,470.2 (obs)). These results verified that *i*-POPFLEX supports efficient incorporation of ncAAs via sense codon reassignment and enables downstream click chemistry for bioconjugation.

**Figure 7.**
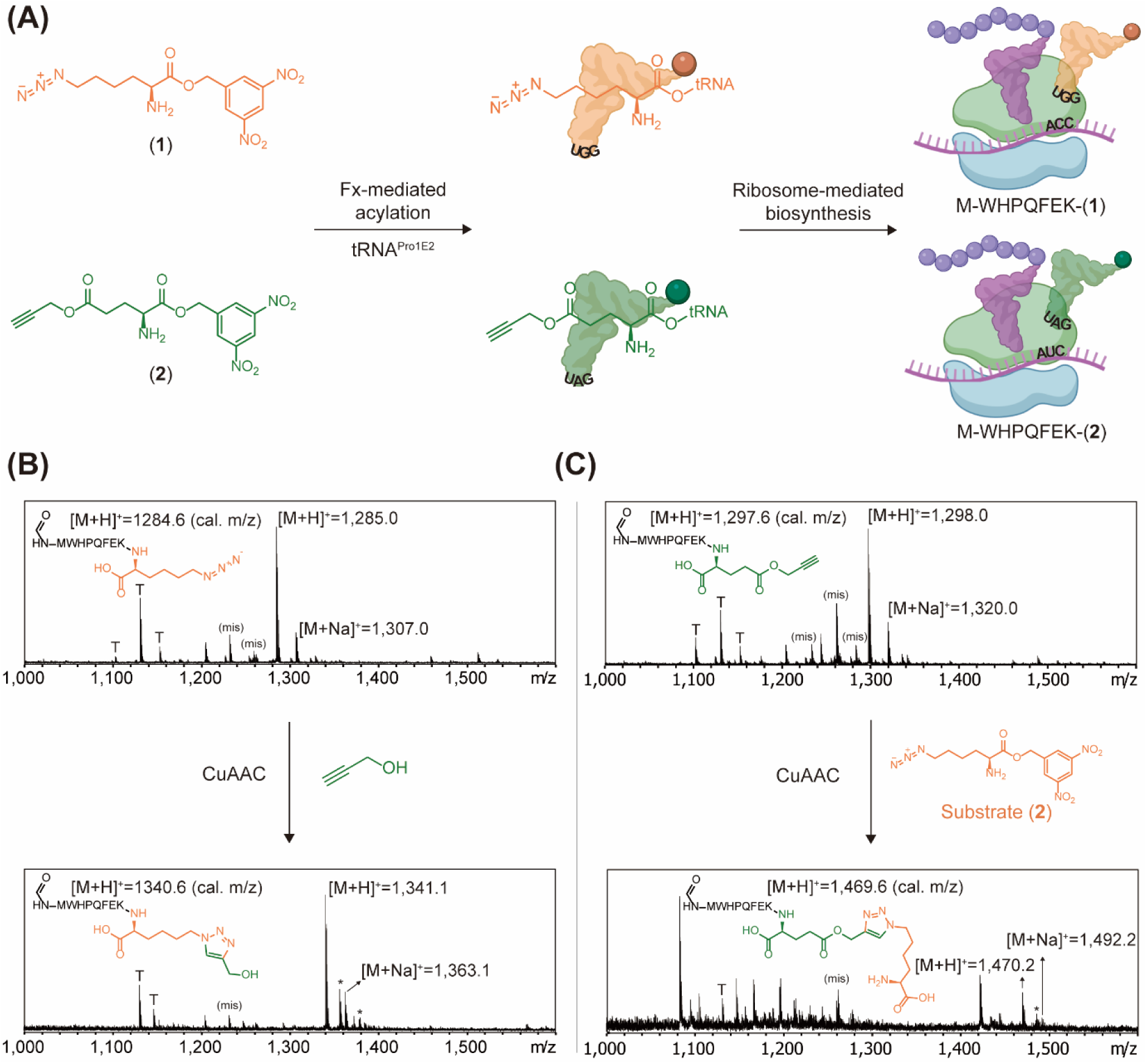
Reducing the aaRS pool minimizes tRNA competition and supports genetic code expansion by reassigning sense codons. (**A**) Synthetic substrates were activated with a DNB ester and individually charged onto tRNAs by Fx. The charged substrate:tRNA conjugates were purified and added to *i*-POPFLEX reaction containing DNA template, enabling incorporation of each substrate through the reassigned sense codon. (**B,C**) MALDI-TOF mass spectra of the resulting peptides. Panels below show MALDI-TOF mass spectra of peptides with a triazole ring. Peaks marked with **T** indicate a truncated peptide that either lacks the last one ncAA at the C-terminus (MWHPQFEK) and peaks with **(mis)** indicate peptide containing a misincorporated Met at the **Y** residue. Peaks marked as an asterisk (*) correspond to peptide containing 2-oxohistidine resulting from histidine oxidation by sodium ascorbate in CuAAC reaction [65]. Of note, the DNB ester of substrate (**2**) was hydrolyzed during the protein translation process. Mass spectra are representative of *n* = 3 independent experiments. The theoretical masses of target peptide in (**B**) and (**C**) are: [M+H]^+^ = 1,284.6 (cal) and [M+Na]^+^ = 1,306.6 (cal), [M+H]^+^ = 1,297.6 (cal) and [M+Na]^+^ = 1,319.6 (cal), respectively. The theoretical masses of target peptide after the CuAAC click reaction in (**B**) and (**C**) are: [M+H]^+^ = 1,340.6 (cal) and [M+Na]^+^ = 1,362.6 (cal), [M+H]^+^ = 1,469.7 (cal) and [M+Na]^+^ = 1,491.7 (cal), respectively.

## Discussion

This work transforms reconstituted CFS from a specialized technique into a practical, automation-ready tool for routine biofoundry use and industrial applications. By replacing gravity-flow chromatography with automated FPLC, we reduced purification times from several hours to ∼20 min, increased factor recovery, and eliminated batch-to-batch variability. The resulting POPFLEX (*in vivo*-produced) mixtures yielded comparable sfGFP fluorescence intensity with the commercial PUREfrex^®^ 2.0 kit, remained fully active after more than two months at −20 °C, and exhibited less than ∼15 % reaction-to-reaction variation on the OT-2 platform (**Figure S1** in the Supplemental Information online). The overall cost for POPFLEX setup was estimated at USD 0.054 µL^-1^ (**Table S1** in the Supplemental Information online), 40 % lower than the previously reported minimum reagent cost (USD 0.09 µL^-1^) [36] and 96 % (25-fold) lower than PURExpress^TM^ (USD 1.36 µL^-1^).

A fully *in vitro* production strategy implemented on the Opentrons OT-2 enabled parallel expression of all 34 translation factors, reducing total process duration from four days to two days. *i*-POPFLEX reactions with commercial ribosomes achieved up to 9.4-fold higher protein yields compared with PURExpress^TM^, but the prohibitive cost (USD 7.74 µL^-1^) limits routine use. To further lower costs, we replaced commercial ribosomes with those harvested directly from an S150 sucrose-cushion pellet (Figure S1 in the Supplemental Information online), thereby eliminating an additional chromatography step [29,65]. Ribosomes obtained from lysate supported lower sfGFP fluorescence than commercial ribosomes, presumably due to the presence of polysomes, which reduce the effective concentration of functional ribosomes. Nevertheless, they still outperformed PURExpress^TM^ (5-fold) and were comparable with PUREfrex^®^ 2.0 (**Figure S1** in the Supplemental Information online). This substitution yielded the significant cost saving, reducing ribosome preparation costs by 3.5-fold (USD 0.02 µL^-1^ vs. USD 0.07 µL^-1^; [36]). In addition, endogenous tRNAs purified from *E. coli* lysate (see Supplemental Information online), reducing reagent costs by a further 9-fold. Overall, the total cost for *i*-POPFLEX setup was USD 0.068 µL^-1^ (**Table S1** in the Supplemental Information online). While this represents a 95 % reduction (20-fold) compared with PURExpress^TM^, it remains higher than POPFLEX because the *in vivo* expression strategy of POPFLEX yields greater protein amounts than the *in vitro* approach used for *i*-POPFLEX. We anticipate that scaling lysate-based reaction and production to higher reaction volume will reduce costs to levels comparable to POPFLEX.

The open architecture of *i*-POPFLEX allows rapid reconfiguration of the translational apparatus and straightforward integration of supplementary chemical and enzymatic reactions. By omitting 12 aaRSs, we created a minimal system that frees several sense codons for reassignment; Fx-charged tRNAs efficiently deliver ncAAs to these codons, and the resulting peptides readily undergo CuAAC.

Looking forward, co-expression of **post-translational modification (PTM)** enzymes [19,66–68], such as disulfide-bond isomerases, glycosyltransferases, kinases, and methyltransferases, could enable cell-free synthesis of proteins or antibodies bearing native PTMs. Combined with ncAA incorporation, this demonstrates that *i*-POPFLEX can produce site-specific bioconjugates and proteins with enzymatic PTMs, expanding applications from ADC assembly to the fabrication of functional biomaterials.

Despite these advances, several challenges remain. First, the workflow still depends on a commercially sourced T7RNAP, which increases costs and may introduce supply-chain risks. Second, the current liquid-handling setup is limited to 48 reactions of 45 µL each per run in 1.5 mL tubes, requiring serial runs or multiple equipment for larger scale and limiting throughput relative to industrial biofoundries. Third, even with the reduced costs, the per-microliter price is still a limiting factor for ultra-large libraries (> 10^9^ variants) unless additional miniaturization or continuous-flow manufacturing strategies are implemented. Finally, advanced strategies for preparing more active ribosomes will be essential to further enhance *i*-POPFLEX performance while lowering costs.

### Concluding remarks

In summary, the speed, cost efficiency, protein synthesis performance, and genetic flexibility achieved by POPFLEX and *i*-POPFLEX position these systems as promising platforms for enzyme screening, synthetic biology prototyping, and emerging applications in biomanufacturing. Further integration with continuous flow factor expression, expanded PTM modules, and machine-learning-guided design cycles could extend cell-free screening to billions of variants and open new opportunities in both fundamental and applied biotechnology (see Outstanding questions).

### Glossary

Aminoacyl-tRNA synthetase (aaRS): An enzyme that catalyzes the attachment of amino acids onto its corresponding tRNA
Cu(I)-catalyzed azide-alkyne cycloaddition (CuAAC): A click chemistry reaction of organic azides and alkynes that synthesizes a triazole ring
Cell-free systems (CFSs): *In vitro* platforms that where biochemical reactions occur
Flexizyme (Fx): An aminoacylating ribozyme that capable of charging a wide range of ncAAs onto tRNAs by recognizing an aromatic group of ester substrates
Elongation factor thermo unstable (EF-Tu): An enzyme that delivers the aminoacyl-tRNA to the A-site of ribosomes
Non-canonical amino acids (ncAAs): Amino acids that are not directly encoded by the natural genetic code
Post-translational modification (PTM): A chemical modification that occurs after protein translation on specific amino acids
RecBCD: A bacterial helicase/nuclease enzyme complex that degrades linear DNA template in the cell-free systems
T7 RNA polymerase (T7RNAP): An enzyme from the T7 bacteriophage that catalyzes RNA synthesis from a DNA sequence with a T7 promoter. T7RNAP is widely used for recombinant protein synthesis.
Transfer RNA (tRNA): RNA molecules that transfer an amino acid to ribosomes

## STAR★METHODS

Detailed methods are provided in the online version of this paper and include the following:

### RESOURCE AVAILABILITY

#### Lead contact

Requests for further information and resources should be directed to and will be fulfilled by the lead contact, Joongoo Lee.

#### Materials availability

All plasmids used in this study are available from the authors upon request.

#### Data and code availability

The data supporting the findings of this study are available in the main text and the supplementary information online. Any additional information required to reanalyze the data reported in this paper is available from the authors upon request.

#### Author contributions

Conceptualization: B.C., Joongoo L.; Data curation: B.C., H.P., H.Jeon, Jihoo L., H.Jin; Investigation: B.C., H.P., H.Jeon, Jihoo L., H.Jin, Joongoo L.; Methodology: B.C., H.P., H.Jeon, Jihoo L., H.Jin, Joongoo L.; Software: B.C., J.H.; Validation: B.C., H.Jeon, Jihoo L.; Visualization: B.C., Joongoo L.; Resources: J.W.L., Joongoo L.; Funding acquisition: Joongoo L.; Project administration: Joongoo L.; Supervision: Joongoo L.; Writing – original draft: B.C.; Writing – review and editing: Joongoo L.

## Acknowledgements

The authors would like to thank Dr. Kosuke Seki (Northwestern University) and Dr. Michael C. Jewett (Stanford University) for providing plasmids for a reconstituted CFS. This research was supported by the Bio&Medical Technology Development Program of the National Research Foundation (NRF) funded by the Korean government (MSIT) (RS-2024-00400486, RS-2025-00520898, RS-2025-16070008, RS-2025-25401663, and RS-2025-23442968). Parts of figure elements were created with Biorender.com.

## STAR★METHODS

### EXPERIMENTAL MODEL AND STUDY PARTICIPANT DETAILS

All culture conditions used in this study can be found in the ‘METHODS DETAILS’ section.

### METHODS DETAILS

#### Construction of DNA templates of POPFLEX components

For One-Pot PURE system, pET and pQE expression plasmids were used for protein expression (**Tables S4** and **S5** in the Supplemental Information online). To establish ssOP system, all genes in pQE expression plasmids were amplified and cloned into a pET21a backbone. Gibson assembly was performed in a 10 µL reaction using the amplified genes and the pET21a vector fragments, with T5 exonuclease, *Taq* ligase, and Phusion DNA polymerase (all purchased from New England Biolabs, NEB). Specifically, 2.5 µL of amplified genes, 2.5 µL of the pET21a backbone, and 5 µL of 2X Gibson Assembly Master Mix were combined and incubated at 50 °C for 50 min. A 2.5 µL aliquot of the assembly mix was used to transform *E. coli* BL21 (DE3) derivative competent cells (NEB).

#### *In vivo* expression of POPFLEX components

The POPFLEX components were prepared using the previously reported methods with modifications [36–38]. Except for EF-Tu, 33 individual *E. coli* strains BL21 (DE3) derivative were transformed with pET plasmids encoding the desired genes and were grown in 150 µL of LB media with 100 µg·mL^-1^ ampicillin in a single 96-well plate sealed with a breathe-easy film (Sigma-Aldrich). Following transformation, cells containing the EF-Tu vector were cultured in a 14 mL round-bottom tube with 3 mL LB media, while the remaining 33 cells were grown in a 96-well plate containing 150 µL LB media per well. After overnight incubation, 1.675 mL of EF-Tu-expressing cells and 55 µL from each of the other cultures were pooled into 500 mL of fresh LB media and grown to saturation overnight at 37 °C with vigorous shaking at 220 RPM in the presence of 100 µg·mL^-1^ ampicillin. Cell growth was monitored, and the protein expression was induced with 0.1 µM (in final) of Isopropyl β-D-1-thiogalactopyranoside (IPTG) when the optical density at 600 nm reached 0.2-0.3. The culture was incubated for an additional 3 h, then harvested by centrifuging at 4,000 g at 4 °C for 10 min. The cells were washed once with pre-chilled PURE buffer A, containing 50 mM HEPES-KOH (pH 7.6), 1 M NH_4_Cl, 10 mM MgCl_2_, and 7 mM 2-mercaptoethanol, and stored at -80 °C. On the following day, the cells were thawed for ∼2-3 h, resuspended in 7.5 mL of PURE buffer A, and lysed using a Q125 Sonicator (Qsonica) with a 3 mm diameter probe at 50 % amplitude using 10 s on/off pulses for a total of 20 min on ice. The cell extract was centrifuged at 21,130 g at 4 °C for 20 min to remove cell debris and was subsequently loaded onto a column for His-tag affinity purification, as described below.

#### Construction of DNA templates for expression of *i*-POPFLEX components

All genes required for protein expressions were amplified and cloned into a pJL1 backbone. Gibson assembly was performed in a 10 µL reaction and a 2.5 µL aliquot of the assembly mix was used to transform *E. coli* DH5-α competent cells (NEB). Recombinant plasmids were extracted using the FavorPrep^TM^ Plasmid DNA Extraction Mini Kit (Favorgen). RF2, methionyl-tRNA formyltransferase (MTF), creatine kinase (CK), myokinase (MK), nucleoside diphosphase kinase (NDK), and inorganic pyrophosphatase (PPiase) were expressed using a linear DNA template, which was prepared by polymerase chain reaction (PCR) and purified via phenol-chloroform DNA extraction. DNA concentration was measured using a Nabi^TM^ Nano Spectrophotometer (MicroDigital Co. Ltd.) at a wavelength of 260 nm.

#### Preparation of *E. coli* S30 cell lysates

*E. coli* S30 cell lysates were prepared using a previously reported method [69]. *E. coli* BL21 (DE3) derivative was grown at 37 °C in 1 L of 2X YTPG media with vigorous shaking at 220 RPM until reaching an optical density at 600 nm ∼0.6, at which point protein expression was induced with 1 mM IPTG. The cells were harvested in the exponential phase (when OD_600_ reached 3.0) by centrifugation at 4,000 g at 4 °C for 10 min, washed three times with pre-chilled buffer A, containing 10 mM Tris-acetate (pH 8.2), 14 mM Mg(OAc)_2_, 60 mM KGlu, and 2 mM dithiothreitol (DTT), and stored at −80 °C. On the following day, the cell pellets were thawed for 2-3 h and resuspended in 1 mL of buffer A per 1 g of wet cell mass. The resuspended cells were lysed using sonication for a total of 2 min on ice, centrifuged two times at 30,000 g, and stored at −80 °C. All buffers described in **Table S8** were freshly prepared before use.

#### *In vitro* expression of *i*-POPFLEX components

Preparation of *i*-POPFLEX components was performed either manually or using an Opentrons OT-2 liquid handler. Each of 34 DNA templates encoding *i*-POPFLEX components was individually mixed with the *E. coli* S30 cell lysates prepared as described above. Specifically, 15 µL of *E. coli* S30 cell lysates, 6.6 µL of Solution A, 6.3 µL of Solution B, and the DNA templates were added to a 1.5 mL microtube to a final volume of 45 µL. Each tube was incubated at 30 °C for 16 h.

For automated *in vitro* expression of *i*-POPFLEX components, an OT-2 protocol was written in Python. *E. coli* S30 cell lysates, Solution A, Solution B, and DNA templates were combined in 1.5 mL microtube to a final volume of 45 µL. Reactions were incubated at 30 °C for 16 h.

After incubation, the reaction mixtures were pooled in a 15 mL conical tube and centrifuged at 21,130 g at 4 °C for 20 min to remove residual cell debris. The supernatant was then mixed with 5 mL of PURE Buffer A and loaded to an ÄKTA system for His-tag affinity purification.

#### Purification of the translational machinery by fast protein liquid chromatography (FPLC)

All His-tagged translation factors were purified through an ÄKTA UPC-900 system (GE Healthcare) using a 5 mL of HisTrap^TM^ HP column (Cytiva) at 4 °C. The column was equilibrated with binding buffer at a flow rate of 3 mL·min^-1^ prior to sample injection. Elution was performed with 450 mM imidazole at the same flow rate, and elution was monitored at 280 nm. Collected fractions were dialyzed three times against PURE dialysis buffer containing 30 % of glycerol for at 4 °C for 1 h. The dialyzed solution was then concentrated using an Amicon^®^ Ultra centrifugal filter (Merck Millipore) with a 3 kDa molecular weight cutoff to a final concentration of 10-12 mg·mL^-1^ for POPFLEX and 5-7 mg·mL^-1^ for *i*-POPFLEX, respectively. The resulting solutions were stored at - 20 °C.

#### SDS-PAGE analysis

For comparison, an additional manual purification was performed to verify the accuracy of the proteins purified by FPLC. Each his-tagged translation factor was purified by using HisPur^TM^ Cobalt Superflow Agarose resins (ThermoFisher Scientific). 30 µL of agarose resins was washed three times with 500 µL of pre-chilled wash buffer (95 % PURE buffer A, 5 % PURE buffer B) in a spin column. The reaction mixture was added to the agarose resins and incubated at 4 °C for 1 h. The resins were washed three times, added 30 µL elution buffer (10 % PURE buffer A, 90 % PURE buffer B), and incubated at 4 °C for 1 h. The supernatant was collected from the column and the proteins were separated via SDS-PAGE. 8 µL of protein samples were mixed with 2 µL of 5X SDS-PAGE Loading Buffer (Biosesang) were boiled at 95 °C for 5 min and electrophoresed on a 12 % SDS-PAGE gel. Separated proteins were stained with Coomassie blue and visualized with a Bio-Rad Gel Doc Go Imaging System.

#### Preparation of in-house energy solution

Energy solution was prepared as described previously with slight modifications [36]. The components of 4X energy solution are 200 mM HEPES-KOH (pH 7.6), 308 mM KGlu, 41.6 mM Mg(OAc)_2_, 8 mM ATP and GTP, 4 mM CTP and UTP, 80 mM creatine phosphate, 8 mM spermidine, 0.08 mM folinic acid, 4 mM TCEP, and 1 mM of each amino acid (**Table S9** in the Supplemental Information online).

#### Purification of ribosome

Ribosomes were purified using a previously reported method [29]. *E. coli* BL21 (DE3) derivative cells were grown at 37 °C in 1 L of 2X YTPG media with vigorous shaking at 220 RPM until an OD600 of 3.0 was reached. Cells were harvested by centrifugation at 4,000 g at 4 °C for 10 min, washed three times with pre-chilled ribosome buffer A, containing 20 mM Tris-HCl (pH 7.2), 100 mM NH_4_Cl, 10 mM MgCl_2_, 0.5 mM EDTA, and 2 mM DTT. Cell pellets were resuspended in 5 mL of ribosome buffer A per gram of wet cell mass and lysed by high-pressure homogenization (VS-4600P, VISON Scientific) at 1,000-1,500 psi. The lysate was centrifuged twice at 30,000 g at 4 °C for 30 min, layered onto a sucrose cushion (1 mL of clarified lysate per 1 mL of ribosome buffer B), and ultracentrifuged at 90,000 g at 4 °C for 18 h. The resulting ribosome pellets were resuspended in ribosome buffer with gentle shaking at 4 °C for 16 h, and the concentration was determined by absorbance at 260 nm, then adjusted to 20 µM. Purified ribosomes were aliquoted, flash-frozen in liquid nitrogen, and stored at −80 °C. All buffers for ribosome purification were freshly prepared and autoclaved before use (**Table S10** in the Supplementary Information online).

#### Purification of *E. coli* total tRNA

*E. coli* total tRNA was purified using a previously reported method with minor modifications [70]. *E. coli* DH5-α was grown in 1 L of LB media to saturation overnight at 37 °C with vigorous shaking at 220 RPM. On the following day, the cells were harvested by centrifugation at 4,000 g at 4 °C for 10 min, washed once with pre-chilled 0.9 % NaCl, and centrifuged again at 4,000 g at 4 °C for 10 min.

For total RNA extraction, the cell pellets were resuspended in pre-chilled tRNA buffer A (50 mM NaOAc, 10 mM Mg(OAc)_2_), mixed with acidic phenol (pH 4.5), incubated at 30 °C for 30 min, and centrifuged at 4,000 g at 4 °C for 10 min. The extraction step was repeated once, and the supernatant was adjusted to a final concentration of 0.2 M NaCl by adding 1.5 mL of 5 M NaCl, followed by isopropanol precipitation (1:1 v/v). The total RNAs were collected by centrifugation at 14,500 g for 25 min at 20 °C.

To remove ribosomal RNA (rRNA), the pellets were washed with pre-chilled 70% ethanol, dried for 10 min, resuspended in 1 M NaCl, and centrifuged at 9,500 g at 4 °C for 20 min. This step was repeated once. The resulting supernatant was mixed with pre-chilled 100 % ethanol (1:2 v/v), incubated at -20 °C for 30 min, centrifuged at 14,500 g for 5 min at 20 °C, and dried for 10 min.

For final precipitation of *E. coli* total tRNA, the supernatant was combined with isopropanol (1.00:0.95 v/v), incubated at -20 °C for 30 min, and centrifuged at 14,500 g at 20 °C for 15 min. The resulting pellets were washed with 70 % ethanol, dried, and dissolved in nuclease-free water.

#### Protein expression on the POPFLEX and *i*-POPFLEX platform

POPFLEX and *i*-POPFLEX reactions were performed using either (A) commercially available Solution I and Solution III from the PUREfrex^®^ 2.0 kit (GeneFrontier) or (B) fully in-house-prepared solution. (A) For reaction with PUREfrex^®^ components, 1 µL of the resulting protein mixture containing the purified 34 proteins (final concentration: 1 mg·mL^-1^) was mixed to a final volume of 5 µL with 2.5 µL of Solution I (buffer, salts, and substrates), 0.5 µL of Solution III (ribosomes), and the DNA template in a 1.5 mL microcentrifuge tube. (B) For fully in-house prepared reactions (5 µL), 1.25 µL of 4X energy solution, 1 µL of protein mixture, 0.25 µL of T7RNAP, 0.5 µL of ribosomes harvested from S150 sucrose-pellet (final concentration: 2 µM), 0.5 µL of purified *E. coli* total tRNA, and DNA template were combined in a 1.5 mL microcentrifuge tube. Fluorescence of sfGFP was measured using a Hidex Sense 425-301 microplate reader, with excitation at 485 nm and emission at 535 nm.

#### Cu(I)-catalyzed azide-alkyne cycloaddition (CuAAC)

CuAAC reaction was carried out as described previously [71]. ncAA-containing peptides were purified using MagStrep “type3” XT beads (IBA) and denatured using 0.1 % SDS in water. For the CuAAC reaction, the purified peptide was combined with 500 µM CuSO_4_, 2.5 mM tris(3-hydroxypropyltriazolylmethyl)amine (THPTA), 1 mM aminoguanidine, and the corresponding reactant; peptides bearing an azide moiety were reacted with an alkyne (prop-2-yn-1-ol), whereas peptides bearing an alkyne moiety were treated with an azide ((*S*)-6-azido-1-((3,5-dinitrobenzyl)oxy)-1-oxohexan-2-aminium; substrate (**1**)). Following incubation at room temperature for 1 h, the resulting peptide was repurified as described above and characterized by MALDI-TOF mass spectrometry.

### QUANTIFICATION AND STATISTICAL ANALYSIS

Each experiment was conducted with at least three replicates. Data are presented as mean ± standard deviation (SD).

### Supplemental information

Document S1. Methods and Materials, Synthesis of ncAAs, Figures S1-S10, Tables S1-S19, Plasmid Map, Linear DNA template sequences, and RNA sequences

Movie S1. *In vitro* expression of *i*-POPFLEX components

Movie S2. Pooling *in vitro*-produced translational machinery

